# Proteogenomics analysis of CUG codon translation in the human pathogen *Candida albicans*

**DOI:** 10.1101/2020.06.03.131292

**Authors:** Stefanie Mühlhausen, Hans Dieter Schmitt, Uwe Plessmann, Peter Mienkus, Pia Sternisek, Thorsten Perl, Michael Weig, Henning Urlaub, Oliver Bader, Martin Kollmar

**Affiliations:** Theoretical Computer Science and Algorithmic Methods Group, Institute of Computer Science, University of Göttingen, Goldschmidtstr. 7, 37077 Göttingen, Germany; Department of Neurobiology, Max-Planck-Institute for Biophysical Chemistry, Am Fassberg 11, 37077 Göttingen, Germany; Bioanalytical Mass Spectrometry, Max-Planck-Institute for Biophysical Chemistry, Am Fassberg 11, 37077 Göttingen, Germany; Bioanalytics Group, Department of Clinical Chemistry, University Medical Center Göttingen, Robert Koch Strasse 40, 37075 Göttingen, Germany; Institute for Medical Microbiology, University Medical Center Göttingen, Kreuzbergring 57, 37075 Göttingen, Germany; Intermediate Care, University Medical Center Göttingen, Robert Koch Strasse 40, 37075 Göttingen, Germany; Group Systems Biology of Motor Proteins, Department of NMR-based Structural Biology, Max-Planck-Institute for Biophysical Chemistry, Am Fassberg 11, 37077 Göttingen, Germany

**Keywords:** Proteogenomics, pathogen, Candida albicans, genetic code

## Abstract

*Candida* yeasts causing human infections are spread across the yeast phylum with *Candida glabrata* being related to *Saccharomyces cerevisiae, Candida krusei* grouping to *Pichia spp.*, and *Candida albicans, Candida parapsilosis* and *Candida tropicalis* belonging to the CTG-clade. The latter lineage contains yeasts with an altered genetic code translating CUG codons as serine using a serine-tRNA with a mutated anticodon. It has been suggested that the CTG-clade CUG codons are mistranslated to a small extent as leucine due to mischarging of the serine-tRNA(CAG). The mistranslation was suggested to result in variable surface proteins explaining fast host adaptation and pathogenicity. Here, we re-assessed this potential mistranslation by high-resolution mass spectrometry-based proteogenomics of multiple CTG-clade yeasts, various *C. albicans* strains, isolated from colonized and from infected human body sites, and *C. albicans* grown in yeast and hyphal forms. Our *in vivo* data do not support CUG codon mistranslation by leucine. Instead, (i) CUG codons are mistranslated only to the extent of ribosomal mistranslation with no preference for specific amino acids, (ii) CUG codons are as unambiguous (or ambiguous) as the related CUU leucine and UCC serine codons, (iii) tRNA anticodon loop variation across the CTG-clade yeasts does not result in any difference of the mistranslation level, and (iv) CUG codon unambiguity is independent of *C. albicans*’ strain pathogenicity or growth form.

## Introduction

*Candida* species belong to the most common fungal pathogens. About 90% of *Candida* infections are caused by just five *Candida* species: *Candida albicans, Candida parapsilosis, Candida tropicalis, Candida krusei* (synonyms *Issatchenkia orientalis*, or lately *Pichia kudriavzevii* (Douglass et al. 2018)), and *Candida glabrata*. The first three of these belong to the so-called CTG-clade, which is termed as such because it only comprises species that have reassigned the CUG codon from leucine to serine (Fitzpatrick et al. 2006). However, CTG codon reassignment is not unique to CTG-clade species but is instead also observed in the *Ascoidea*-clade and in a lineage comprising *Pachysolen* and *Nakazawaea* yeasts (Mühlhausen et al. 2016, 2018; Krassowski et al. 2018). While the CUG codon is translated as serine in most *Ascoidea*-clade species, *Ascoidea asiatica* exceptionally translates CUG stochastically into both serine and leucine (Mühlhausen et al. 2018). In *Pachysolen* and *Nakazawaea* species CUG is translated into alanine (Mühlhausen et al. 2016, 2018). The CUG decoding 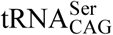 in the CTG-clade originated from a 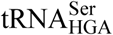 isoacceptor (for UCU, UCG and UCA codons), whereas the 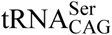 in *Ascoidea*-species originated from a 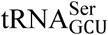 (for AGY codons) (Mühlhausen et al. 2018). Accordingly, the long-favoured “ambiguous intermediate” model for CTG reassignment became extremely unlikely as it would require multiple independent events including divergence of multiple different tRNA-types as ambiguous intermediates. Currently, the best model to explain the reassignments is the “tRNA-loss driven codon reassignment” hypothesis (Mühlhausen et al. 2016; Kollmar and Mühlhausen 2017b). According to this model, 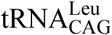 was lost in the last common ancestor of Ascoideae, Pichiaceae, Saccharomycetaceae, and CTG-clade yeasts. The then unassigned CUG codon was captured by leucine-, serine-, and alanine-tRNAs which are incidentally the only tRNA-types where anticodons are not part of tRNA identity elements.

A few CTG-clade species such as *C. albicans* and *Candida maltosa* have been reported to translate the reassigned CTG codon by 3-5% into leucine (Suzuki et al. 1997; Gomes et al. 2007). Central to *C. albicans*’ pathogenicity is the ability to change the cellular morphology between the yeast and mycelial forms (Tsui et al. 2016). In this process stochasticity of cell surface proteins might increase *C. albicans*’ ability to host adaptation (Miranda et al. 2013). In contrast, other species including *Candida cylindracea* have been reported to translate CUG unambiguously (Suzuki et al. 1997). Translation ambiguity was proposed to depend on tRNA sequence. In particular, a guanosine at position 33, 5’-adjacent to the CAG anticodon (G33), and a 1-methyl guanosine nucleotide at position 37, 3’-adjacent to the anticodon (m^1^G37), were identified to be invariant in all 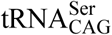. The only exception was the *Candida cylindracea* tRNA in which an adenosine is found at position 37 (A37) (Yokogawa et al. 1992; Suzuki et al. 1994). While the m^1^G37 mutation was shown to cause an increasing leucylation level, the G33 mutation prevented LeuRS (leucine-tRNA synthetase) binding. This suggested a balancing effect of the two mutations leading to low level mischarging of CTG-clade 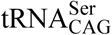 with leucine (Santos et al. 1997). Incorporation of 3-5% of leucine at CUG were reported from a genetic rescue experiment (Suzuki et al. 1997) and mass spectrometry of an overexpressed peptide (Gomes et al. 2007). In contrast, very low levels of mistranslation by leucine (1.45 ± 0.85% in a *C. albicans* control and 0.64 ± 0.82% in a knock-out of one of the two 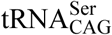) were found in fluorescence measurements (Bezerra et al. 2013). Similarly, we observed CUG translation into leucine only at background ribosomal mistranslation rates in high-resolution proteogenomics experiments in *Clavispora lusitaniae* and *Babjeviella inositovora* (Mühlhausen et al. 2018). These results prompted us to reinvestigate CUG codon translation across the CTG-clade performing state-of-the-art proteogenomics analyses.

## Results

### Selection of yeasts for analysis

To determine the accuracy of CUG codon translation dependent on tRNA identity elements *in vivo* across the CTG-clade we aligned 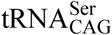 from *C. cylindracea* and 38 sequenced species (Figure 1) (Mühlhausen et al. 2018). The 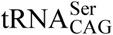 most similar to that of *C. cylindracea* is *B. inositovora.* Both have an A37 instead of the common m^1^G37, a substitution which has been suggested to suppress mischarging by the LeuRS. This might explain the observed unambiguous translation of CUG as serine in *B. inositovora* (Mühlhausen et al. 2018). *C. lusitaniae’s* 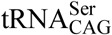 is as diverged from *C. albicans* 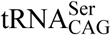 as *B. inositovora* and *C. cylindracea* 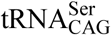 are from the *C. albicans* 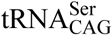, but the differences compared to the *C. albicans* 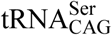 are at non-overlapping positions (Figure 1). Anticodon loops are identical between *C. albicans* and *C. lusitaniae* 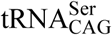, thus some of the 21 nucleotide differences spread across the other loops must explain why CUG is translated unambiguously in *C. lusitaniae* but not in *C. albicans* (Mühlhausen et al. 2018). To best represent 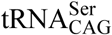 diversity seen across the CTG-clade we selected *Candida tropicalis, Millerozyma acaciae, Candida dubliniensis* and *C. albicans* for an in-depth analysis. *C. tropicalis* 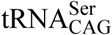 and the identical *Candida sojae* 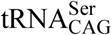 are the only 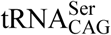 with a cytosine at position 33 (Figure 1). Substitution of the conserved G33 by cytosine has been shown to strongly enhance leucylation activity *in vitro* (Suzuki et al. 1997). *M. acaciae* 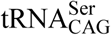 differs from *C. albicans* 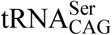 only in ^e1^U^e2^C at the tip of the variable loop. This sequence is found in most CTG-clade yeast 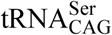 (Figure 1). Corresponding nucleotides in *C. albicans’* 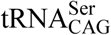 are ^e1^A^e2^U. To determine whether other factors than 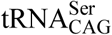 sequence could influence CUG translation ambiguity we selected *C. dubliniensis*, whose 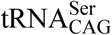 is identical to that of *C. albicans*, and we choose multiple different *C. albicans* strains. In addition, we analysed *C. albicans* in yeast and hyphal growth forms.

**Figure 1.**
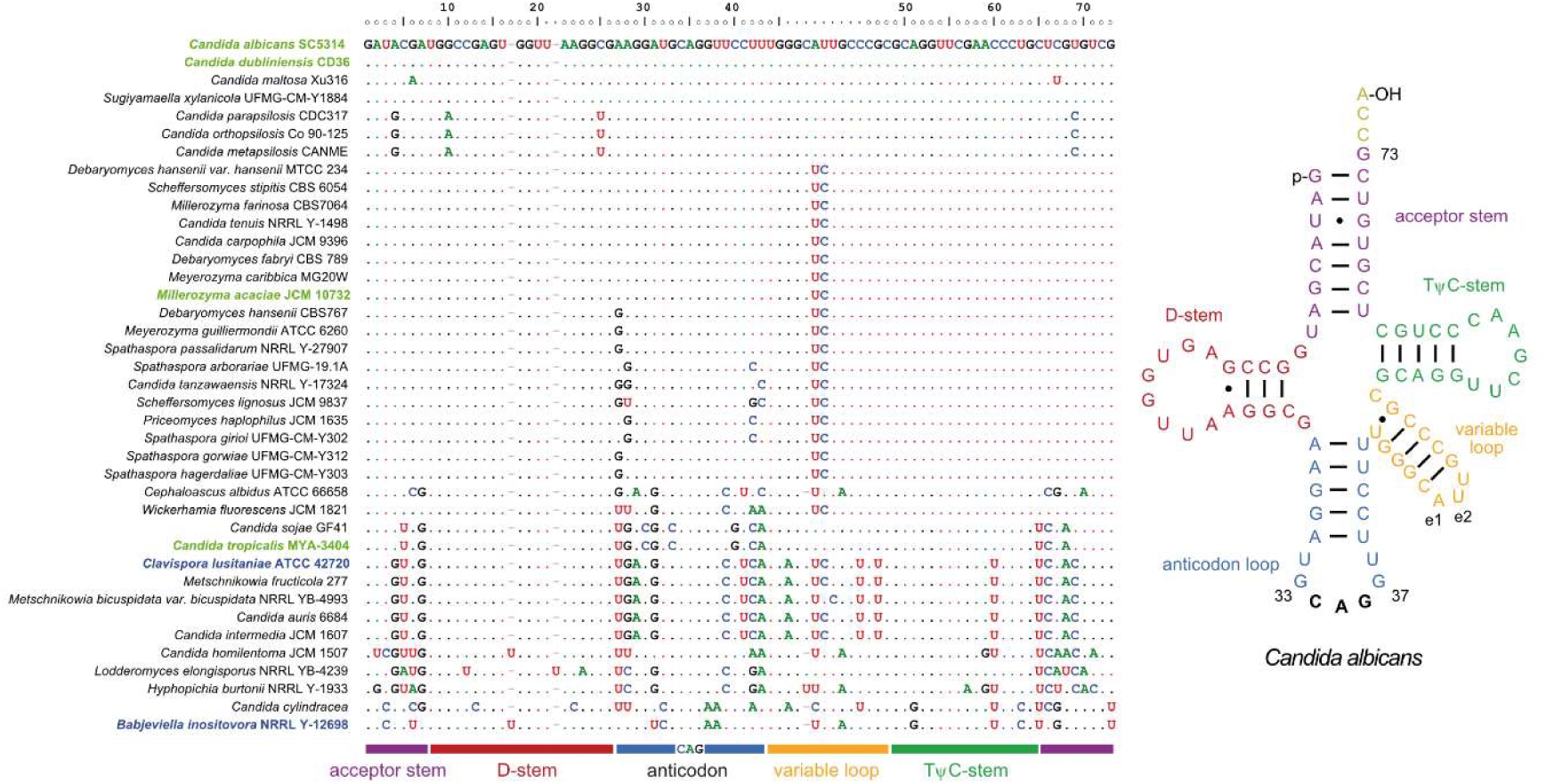
Serine-tRNA(CAG) diversity in CTG-clade yeasts. Alignment of 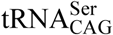 from 38 sequenced yeast genomes and *Candida cylindracea*. Nucleotides identical to the *Candida albicans* 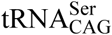 are represented by dots to better highlight divergence. Numbering of nucleotide sequence positions according to standard nuclear tRNA numbering (Sprinzl et al. 1998). Species with high-resolution proteomics data available from previous studies (Mühlhausen et al. 2018) highlighted in blue, species analysed in this study highlighted in green.

CTG-clade 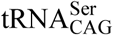 are undoubtedly Ser-tRNAs (Mühlhausen et al. 2016, 2018) but differ within the group in up to 25% of their nucleotides. They all have the Leu-tRNA CAG anticodon triplet and most have m^1^G37, which is present in most but not all Leu-tRNAs. Based on these few identities it seems exaggerated (albeit commonly done) to term the CTG-clade 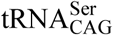 ‘chimeric’ tRNAs. A more concise use of ‘chimera’ would encompass only those entities consisting of clearly distinct parts of independent origins with each having a substantial impact on the entity. The 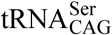 originated from a 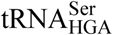 isoacceptor followed by single nucleotide insertion and/or point mutation and not by joining pieces from different tRNA genes (Mühlhausen et al. 2018).

### Unbiased peptide spectrum matching with database replicates

To determine the CUG codon translation in the selected yeasts, we performed liquid chromatography-tandem mass spectrometry (LC-MS/MS) analysis of cell lysates (Supplementary Table S1). We did not employ a de novo peptide sequencing approach as this is not accurate enough to evaluate spectra of rare peptides (Devabhaktuni and Elias 2016; Muth and Renard 2018). This approach mistakenly suggested 10% of CUGs mistranslated by leucine (Riley et al. 2016) instead of the unambiguous translation into alanine in *Pachysolen tannophilus* as reported elsewhere (Mühlhausen et al. 2016). In contrast, database searches with unbiased databases allow precise detection of rare peptides with a precision of single amino acid differences being mapped to identical protein regions, which can result from low-level ribosomal mistranslation (Mühlhausen et al. 2016) or stochastic translation as in *Ascoidea asiatica* (Mühlhausen et al. 2018). Thus, we followed our previously described approach and generated an unbiased database for each analysed species containing each CUG codon (or, respectively, another codon of interest) translated into each of the 19 amino acids (leucine and isoleucine are indistinguishable by MS/MS and thus translation into isoleucine was omitted). The database search algorithm has no a priori knowledge about the “correct” translation and therefore treats all spectrum matches equivalent during scoring and filtering. In addition to applying a false discovery rate (FDR) of 1% for global quality filtering, we further filtered spectrum matches of peptides containing CUG-translated amino acids by requiring residues at CUG positions to be supported by b- and/or y-type fragment ions on both sides. The majority of these amino acids, 89 to 95% depending on sample, are part of extended chains of b-/y-type supported amino acids (Supplementary Table S1).

### Unambiguous translation of CUG as serine in all CTG-clade yeasts

Analysing 11,084,524 MS/MS spectra using this approach, we recovered 10 to 18% (5 to 12% with b-/y-type support) of the total CTG positions in the genomes, spread across 1,000 to 2,400 genes (Table 1). These values are similar to those obtained from the previously analysed yeasts *B. inositovora* and *C. lusitaniae*. CTG position coverage correlated with the number of measured and processed spectra, which is expected given very similar numbers of CTG codons in the genomes of *C. albicans, C. dubliniensis*, and *C. tropicalis* as compared to 1.5 and 2-fold more in the genomes of *B. inositovora* and *C. lusitaniae*, respectively. When comparing observed CUG codon translations we found 99% ± 1% (on average 485 positions, numbers from here on refer to b-/y-type supported positions) CUG codon positions in *C. albicans* to be translated unambiguously, namely 95% (average of 464 positions) translated into serine and 4% (average of 21 positions) into any other amino acid (including, but not exclusively, leucine or isoleucine; Figure 2). The CUG codons translated as leucine or isoleucine are most affected by processing the data against diploid (SC5314 strain) versus pseudo-haploid (WO-1 strain) annotations indicating strain and/or allele differences and not mistranslations (more details below). At the remaining 1% (average 4 positions) CUG positions, peptides with at least two different CUG codon translations were found. The unambiguous serine-translated CUG codon positions were supported by an average of 1961 PSMs per sample (on average 4 PSMs per position; PSM = peptide spectrum match). At ambiguous positions, 12 times more PSMs with CUG translated as serine compared to all other amino acids were found (Figure 2). Notably, only an average of 4 (0.2%) PSMs were found with CUG codons translated as leucine or isoleucine at ambiguous positions (average one position, or 0.2%). In *C. dubliniensis* 94% of the PSMs mapped to unambiguous positions and only a single CUG position with ambiguous translation was found. Twenty-seven PSMs contained this position translated as serine and one PSM as aspartate. Similarly, in *M. acaciae* no CUG codon position ambiguously translated with leucine or isoleucine was found. In *C. tropicalis* 40 (2.2 %) of the 1827 PSMs mapped peptides with CUG codons translated as leucine or isoleucine. However, eleven of these mapped to codons where no other translation than leucine or isoleucine was found, and the other 29 mapped to a single CUG codon position in the ubiquilin gene, which is involved in protein degradation. At the same codon position a minority of nine PSMs had the CUG translated as serine. Although 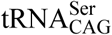 of most of the analyzed yeasts differ from the *C. albicans* 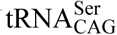, these *in vivo* data show that individual sequence differences do not correlate with and result in a supposed CUG mistranslation into leucine/isoleucine. *C. dubliniensis* with an identical 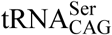 compared to *C. albicans* did not show any ambiguity towards leucine.

**Table 1.** Number of obtained PSMs and proteins and CTG positions covered. If not indicated otherwise, numbers include positions both with and without direct b-/y-type support.

| Yeast and protein database | Strain | Growth form | # PSM with CTG | # PSMs with support ed CTG | # CTG pos covered | # support ed CTG pos covered | % support ed pos | # identifie d proteins with CTG | % identifie d proteins with CTG | CTG pos recovery [%] |
| --- | --- | --- | --- | --- | --- | --- | --- | --- | --- | --- |
| *Candida albicans* SC5314 | DSM70014 | yeast | 3510 | 2206 | 345 | 243 | 70.43 | 1406 | 17.46 | 9.85 |
| *Candida albicans* SC5314 | SC5314 | yeast | 1761 | 1024 | 658 | 413 | 62.77 | 2151 | 26.71 | 10.69 |
| *Candida albicans* SC5314 |  | hyphal | 6368 | 3549 | 1148 | 719 | 62.63 | 2687 | 33.37 | 14.1 |
| *Candida albicans* SC5314 | EU0006 | yeast | 3801 | 2078 | 762 | 469 | 61.55 | 2306 | 28.64 | 11.3 |
| *Candida albicans* SC5314 |  | hyphal | 4021 | 2365 | 934 | 608 | 65.1 | 2503 | 31.09 | 12.38 |
| *Candida albicans* SC5314 | EU0009 | yeast | 4838 | 2754 | 931 | 599 | 64.34 | 2477 | 30.76 | 12.59 |
| *Candida albicans* SC5314 |  | hyphal | 4405 | 2467 | 925 | 610 | 65.95 | 2515 | 31.23 | 12.04 |
| *Candida albicans* SC5314 | EU0075 | yeast | 3284 | 1611 | 756 | 413 | 54.63 | 2240 | 27.82 | 11.47 |
| *Candida albicans* SC5314 |  | hyphal | 3501 | 1762 | 876 | 499 | 56.96 | 2446 | 30.38 | 11.7 |
| *Candida albicans* WO-1 | DSM70014 | yeast | 3195 | 2048 | 329 | 223 | 67.78 | 1197 | 30.27 | 11.63 |
| *Candida albicans* WO-1 | SC5314 | yeast | 1501 | 855 | 592 | 371 | 62.67 | 1756 | 44.41 | 12.47 |
| *Candida albicans* WO-1 |  | hyphal | 5414 | 3015 | 1003 | 644 | 64.21 | 2136 | 54.02 | 16.25 |
| *Candida albicans* WO-1 | EU0006 | yeast | 3436 | 1862 | 719 | 435 | 60.5 | 1861 | 47.07 | 13.89 |
| *Candida albicans* WO-1 |  | hyphal | 3619 | 2093 | 888 | 572 | 64.41 | 1999 | 50.56 | 15.59 |
| *Candida albicans* WO-1 | EU0009 | yeast | 4412 | 2423 | 882 | 557 | 63.15 | 1981 | 50.1 | 15.67 |
| *Candida albicans* WO-1 |  | hyphal | 4066 | 2217 | 872 | 570 | 65.37 | 2005 | 50.71 | 15.06 |
| *Candida albicans* WO-1 | EU0075 | yeast | 2957 | 1424 | 709 | 381 | 53.74 | 1795 | 45.4 | 14.24 |
| *Candida albicans* WO-1 |  | hyphal | 3166 | 1568 | 834 | 471 | 56.47 | 1959 | 49.54 | 14.79 |
| *Candida dubliniensis* | CBS 7987 | yeast | 966 | 466 | 232 | 121 | 52.16 | 1003 | 25.61 | 10.02 |
| *Candida tropicalis* | DSM 24507 | yeast | 3032 | 1827 | 378 | 262 | 69.31 | 1247 | 31.45 | 12.64 |
| *Millerozyma acaciae* | CBS 5656 | yeast | 4962 | 2895 | 1631 | 1111 | 68.12 | 2418 | 67.62 | 17.85 |
| *Babjeviella inositovora* | NRRL Y-12698 | yeast | 4937 | 2348 | 1211 | 930 | 76.8 | 2122 | 47.47 | 17.34 |
| *Clavispora lusitanae* | ATCC 42720 | yeast | 5803 | 2715 | 2360 | 1835 | 77.75 | 2843 | 56.95 | 15.87 |

**Figure 2.**
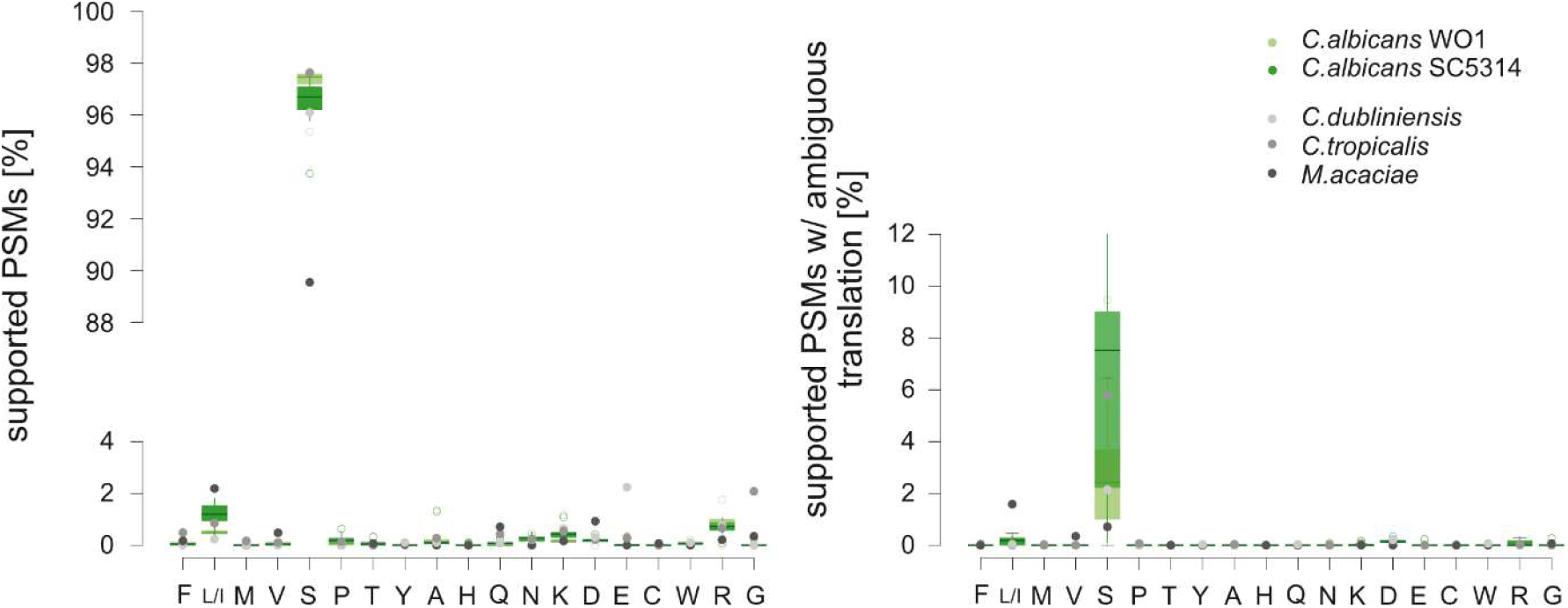
Percentage of PSMs containing a b-/y-type fragment ion supported CUG codon by CUG translation. Values for *C. albicans* samples are depicted in green boxplots (light green: samples searched against a database derived from WO-1 genome annotation; dark green: samples searched against a database derived from SC5314 genome annotation). Values for *C. dubliniensis, C. tropicalis* and *M. acaciae* samples added for comparison. Values for any identified, supported CUG position (left side) are contrasted on the right with values for those CUG positions at which multiple CUG codon translations have been found. The prevalent CUG translation, serine, is found in most of the corresponding PSMs. The numbers of CUG translations other than serine is higher for the allele-resolved *C. albicans* SC5314 database analyses than for the pseudo-haploid WO-1 database analyses indicating strain differences and haplotype merging effects. (Mis-)translation into leucine/isoleucine is, contrary to what is expected based on a suspected 3% mischarging of 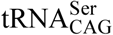 with leucine, no more prevalent than mistranslation into any other amino acid.

### Amino acids other than serine at CUG positions

Across the nine *C. albicans* samples (five strains, four of these each in yeast and hyphal growth forms), 423 CUG codon positions were found with the CUG codons translated by other amino acids than serine (Figures 2 and 3A). These translations could be the result from strain differences (search databases were generated from the SC5314 and WO-1 strains), allele-specific expression (the WO-1 is a pseudo-haploid genome assembly) or codon mistranslation while the correctly translated peptide is missing. Strain differences and allelic expression should be visible in multiple samples, while mistranslated peptides as result from a random process should be unique. At 165 (39.0%) CUG codon positions, the identical translation was found in at least two different samples indicating strain or allelic expression differences. At the other 258 CUG positions, peptides found were unique to one of the nine samples. However, only 7.0% of these peptides had leucine (or isoleucine) at the CUG position implying that if these were the result of mistranslation (e.g. from leucine-mischarged 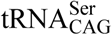), mistranslation by other amino acids must be more prevalent.

**Figure 3.**
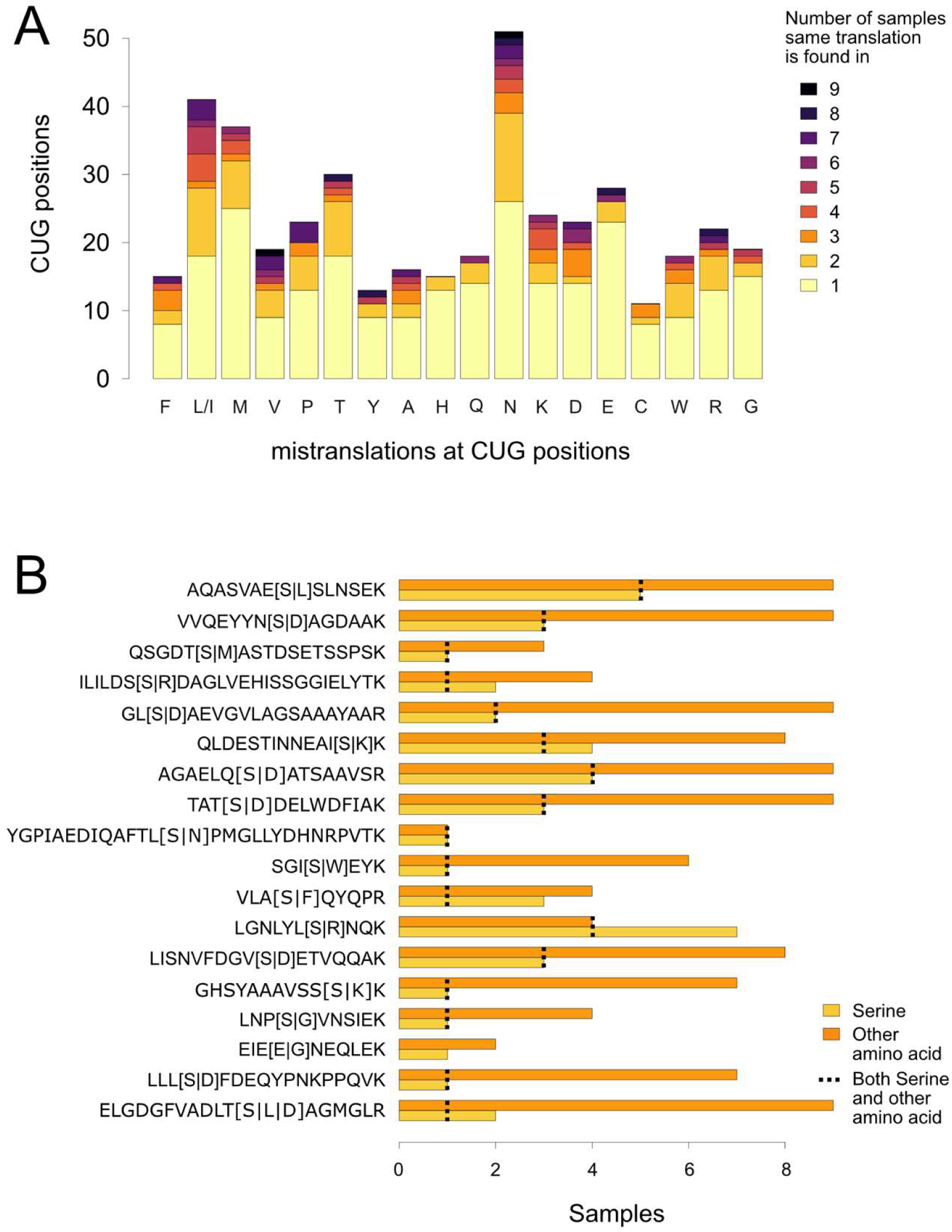
CUG codon positions with translations other than serine. A) The number of CUG codon positions at which only amino acids other than serine were found separated by their identification in only one or multiple of the nine samples (five *C. albicans* strains, yeast and hyphal growth forms). B) The plot lists the eighteen CUG codon positions for which peptides with different CUG translations were found. Translations are distinguished into “serine” and “any other amino acid”. Alternative translations are indicated in brackets in the peptide sequences. The number of samples in which multiple translations were found in a single sample are indicated with a dotted line. For peptides [1] and [3] (read from the bottom) different non-serine translations were found in different samples.

Across all nine samples, at together 18 CUG positions peptides with the CUG translated into different amino acids were found (Figure 3B). At ten of these positions, the combination of serine and another amino acid (leucine or isoleucine in one case) was found in at least two samples indicating allelic expression. At six positions, ambiguous translation with serine and another amino acid (but not leucine or isoleucine) was found in a single sample, at one position, glutamate and glycine but no serine were found, and at another position three amino acids, serine, aspartate and leucine or isoleucine, were found. These cases indicate that only at a single CUG position across nine samples an ambiguous leucine (or isoleucine) was found, which rather demonstrates low level mistranslation than CUG ambiguity.

On average, 2073 (0.86%) of the PSMs of each *C. albicans* sample span a CUG codon. If a 3% mischarging of the 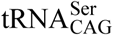 by leucine was assumed as proposed by earlier studies (Suzuki et al. 1997; Gomes et al. 2007), an average of 62 PSMs with the CUG codon translated by leucine are to be expected per sample. This is in sharp contrast to the experimentally found average of only 17 PSMs that further separate into those unique for the position (no serine translation at the position found) and those at the two positions where serine and leucine or isoleucine were found at the same time.

### Level of ambiguous translation (mistranslation) is below bacterial contamination

To obtain a reference for peptide detection levels using our approach, we mixed *M. acaciae* with *E. coli*. As starting point a sample of 90% *M. acaciae* cell lysate and 10% *E. coli* cell lysate was prepared. This sample was diluted by half with *M. acaciae* cell lysate in several steps to a final concentration of 0.16% *E. coli* (Figure 4). At this low concentration, *E. coli* proteins could still be well detected. For comparison, we analysed the supposedly “*E. coli* free” *M. acaciae* sample and several *C. albicans* samples combining the yeast-specific databases with the *E. coli* database. This analysis revealed a considerable fraction of *E. coli* contamination in all samples although highest standards were followed for clean work (Figure 4B). The total numbers of PSMs based on translated CUG codons in all samples correlate well with the percentage of *E. coli* added to the *M. acaciae* sample (compare Table 1 and Figure 4A). Accordingly, the observed fraction of CUG codons translated by other amino acids than serine is clearly below the level of *E. coli* contamination in the samples.

**Figure 4.**
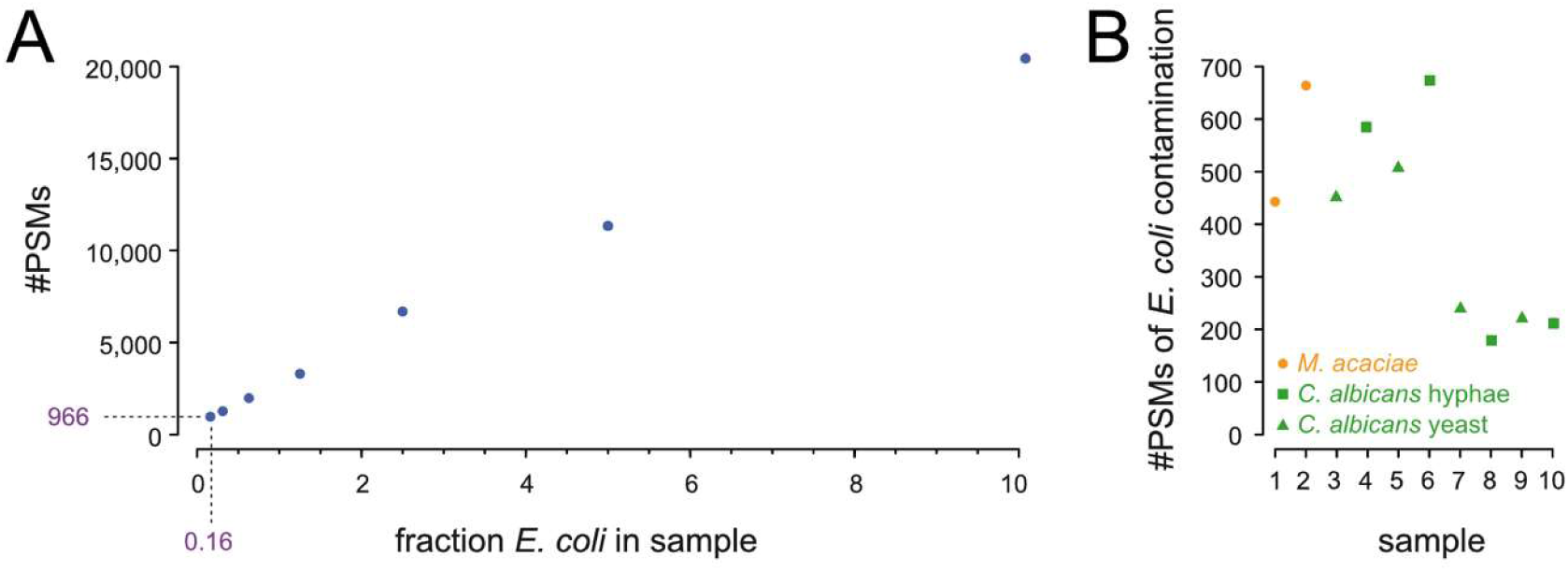
Analysis of *E. coli* contamination level. A) Number of PSMs matching *E. coli* proteins in an otherwise *M. acaciae* sample by fraction of *E. coli* in the sample. B) Number of *E. coli* PSMs found in samples that where not purposely mixed with *E. coli*. Samples 1 and 2 are *M. acaciae* samples used as control to determine the *E. coli* contamination level in a supposed “*E. coli* free” sample. Samples 3 to 8 are *C. albicans* strains SC5314, EU0006, EU0009 and EU0075, each in yeast and hyphal growth form. The number PSMs found for background *E. coli* contamination is below the mixture sample with 0.16% *E. coli* and 99.84% *M. acaciae*. The number of *E. coli* PSMs was normalized to 100,000 PSMs for each sample.

### Are other codons as unambiguous as the CUG codon?

With a codon usage of 0.42%, CTG belongs to the rarely used codons in the *C. albicans* genome annotation (Skrzypek et al. 2017). To determine whether the observed unambiguous CUG codon translation depends on the global codon frequency and/or the type of amino acid, we analysed the CUU leucine and the UCC serine codons, which are used with frequencies of 0.28% and 0.87% in the genome, respectively. Although their global genomic codon usage differs by a factor of three as does the number of PSMs covering the respective codons, the number of CUU and UCC codons covered with PSMs was almost identical (Supplementary Tables S2-S4). Strikingly, about four to five times more CUU and UCC codons than CUG codons were found to be covered by PSMs with the respective position(s) being supported by b-/y-type fragment ions. This indicates that CUU and UCC codons are present in commonly expressed genes while the CUG codons are rather enriched in genes with low or no detectable expression level. Similar to CUG, both CUU and UCC codons show a low level of ambiguous translation with 0.73% and 5.09% of codon positions being covered by peptides with at least two different translations (Figure 5 left). On average, 2.16% and 22.01% of PSMs covering the CUU and UCC codons translated as leucine and serine, respectively, were found at positions where PSMs with other translations of the codon were also observed showing dominance of the standard translation (Figure 5 right). Together, this indicates that the UCC serine codon tolerates a slightly higher level of mistranslation than the CUU leucine codon, likely because serine amino acid positions are enriched at the surface of proteins and are generally less conserved than leucine amino acid positions (Mühlhausen and Kollmar 2014; Mühlhausen et al. 2018). This analysis of other, related codons demonstrates that the CUG codon is translated in *C. albicans* as unambiguously as other codons. If CUG was supposed to be translated ambiguously, a similar ambiguity would have to be assumed for other codons as well.

**Figure 5.**
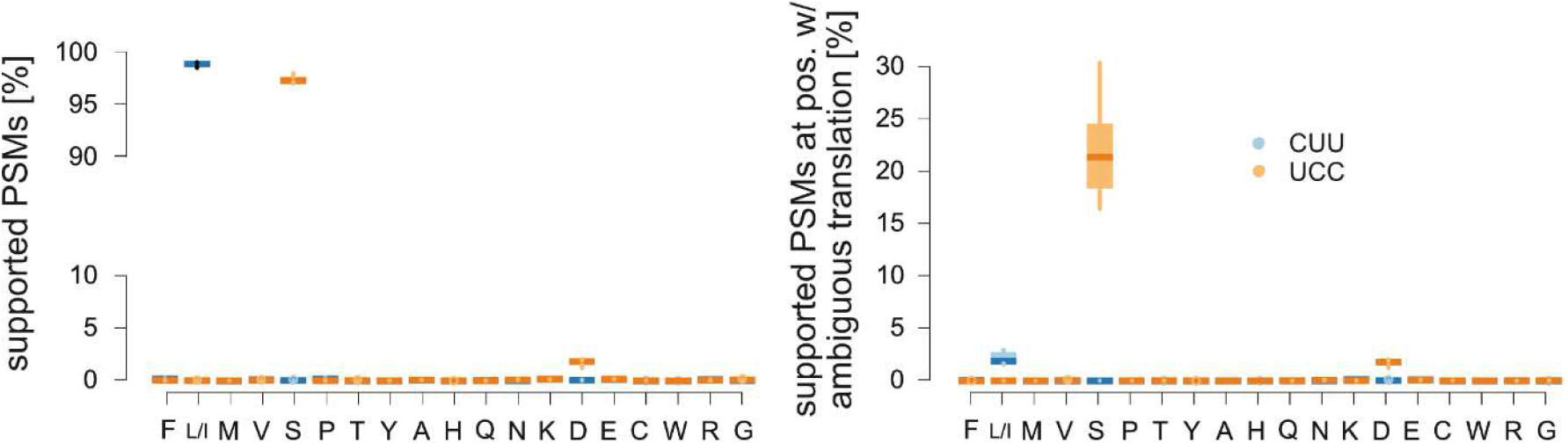
Percentage of PSMs containing a b-/y-type fragment ion supported CUU (leucine; blue) or UCC (serine; orange) codon by translation. *C. albicans* samples have been searched against CUU and UCC databases derived from WO-1 genome annotation. Values for any CUU (UCC) codon position covered with PSMs on the left are contrasted with those for positions that have been identified as being ambiguously translated on the right. Far more UCC positions than CUU positions are translated ambiguously.

### Unambiguous translation of CUG as serine in *C. albicans* yeast and hyphal growth forms

To investigate whether the reported CUG ambiguity is strain or growth form dependent, we tested five strains, DSM70014, SC5314 (clade 1), and three genetically distinct clinical isolates EU0006 (clade 2), EU0009 (clade 12), and EU0075 (clade 4). The latter four strains were grown in yeast and hyphal growth form. All analyses resulted in the same unambiguity of the CUG translation (Table 1, Figure 1).

## Discussion

Our unbiased, statistical evaluation of proteome data shows that CTG-clade yeasts do not translate the CUG codon ambiguously *in vivo*. Although there is a measurable level of CUG mistranslation, it is (i) similar to that of the leucine CUC and serine UCC codons, (ii) similar in six CTG-clade species covering different substitutions of the conserved guanosine nucleotides adjacent to the CAG anticodon in 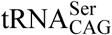, and (iii) similar in yeast and hyphal growth forms of *C. albicans*. Last but not least, mistranslation of CUG into leucine (or isoleucine) is not preferred, although this would be expected if the 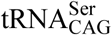 were partially mis-charged by leucine. Instead, several other amino acids were found at similar levels at CUG codon positions.

How do our findings relate to previous reports of a slight CUG mistranslation into leucine? In 1997, tRNA pools were purified from *C. cylindracea* and *Candida zeylanoides* and radioactively labelled immediately (Suzuki et al. 1997). The 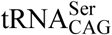 were subsequently pulled out of the mixture using a solid-phase attached DNA probe. This last step is crucial because the DNA probe could be selective enough to extract the *C. cylindracea* 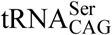 but not to quantitatively exclude all *C. zeylanoides* Leu-tRNA. The entire approach allows for accumulating contaminations at multiple steps and control experiments using other tRNAs are missing. In the same study, the authors performed a genetic rescue experiment introducing a plasmid encoding the *Saccharomyces cerevisiae URA3* gene, which contains a leucine essential for activity, into *Candida maltosa* (Suzuki et al. 1997). While *C. maltosa* was not viable with a serine codon at the essential leucine position, weak growth was observed in case of a CUG codon. However, experiments such as this performed under strong selection do not allow distinguishing translation by mis-charged 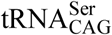 from mistranslation by non-cognate Leu-tRNAs. The latter argument might also explain the observed incorporation of leucine into a reporter peptide *in vivo* (Gomes et al. 2007). In that study, a peptide containing a serine decoded by a CUG codon was fused to a reporter protein, the protein overexpressed in *C. albicans*, then purified and in-gel digested, and the resulting peptides identified and quantified using high-pressure liquid chromatography and tandem mass spectrometry. As control, the authors analysed potential mistranslation of lysine AAA and aspartate GAU codons by near-cognate Asn- and Glu-tRNAs, respectively (Gomes et al. 2007). However, yeast genomes only contain the near-cognate 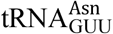 and 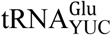, which do not allow translation of AAA and GAU codons by standard or wobble base pairing rules (Crick 1966; Kollmar and Mühlhausen 2017a). Therefore, such mistranslations were, in fact, not observed. In contrast, the serine CUG codon can be mis-translated by the non-cognate 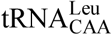 establishing a C•A mispair at the first codon-anticodon position (Rozov et al. 2015). It is also known that stress increases mistranslation levels in general (Wang and Pan 2016; Mohler and Ibba 2017) and likely this is what the authors observed when co-expressing a *S. cerevisiae* mutant 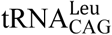 (Gomes et al. 2007). Unfortunately, control experiments using the identical peptide but testing other codons for potential mistranslations caused by wobble mis-pairings were not performed there. Also, mistranslation by leucine versus mistranslation by isoleucine cannot be distinguished.

It could be possible that we did not observe CUG codons mistranslated as leucine because they were not in the prepared cell lysates. To considerable extent CUG codons in *C. albicans* and also the other CTG-clade yeasts are found at conserved serine positions in proteins (Mühlhausen and Kollmar 2014; Mühlhausen et al. 2018) but never at conserved leucine positions. Thus, proteins with leucines randomly introduced at important serine positions could immediately be degraded. However, *C. albicans* with more than 25% of the CUG codons artificially translated as leucines was shown to be viable (Gomes et al. 2007; Bezerra et al. 2013). Alternatively, CUGs translated to leucine could be enriched in cell wall proteins that were not present in the cell lysate. However, we are not aware of any mechanism that would preferentially select Ser-tRNAs charged with leucine (as suggested by the 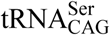 mischarging hypothesis) for synthesizing cell wall and secretome proteins while excluding these mischarged 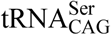 for soluble proteins. Also, we are not aware of any mechanism by which ribosomes could select mischarged 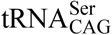 instead of correctly charged 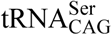 in a sequence-dependent context such as the seleno-cysteine incorporation at stop codons. In the latter case a tRNA is selected instead of a release factor, which are completely different entities compared to correctly charged and mischarged 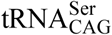, which are identical except for a hydroxyl and an isopropyl group at the amino acid side chain.

Are our proteogenomics experiments sufficiently reliable to determine codon mistranslation? There are several findings that support our conclusions. First, the observed CUG codon ambiguity is below the level of bacterial contamination. We have taken every effort to prepare pure samples and high experimental standards are indeed a requirement for working with *C. albicans*. Nevertheless, *E. coli* proteins could be detected at very low levels. Second, we did find some CUG codon positions with translations other than serine, and even positions with multiple different translations. This ambiguity is either caused by mistranslation or by sequence differences between the alleles. If we assumed no allelic differences or expression of only one of the two alleles in this diploid species, the maximum possible mistranslation level would then correspond to the observed level of other amino acids found at CUG codon positions. However, explaining the non-serine translation of CUG codons by strain and allelic differences is much more likely because most of these translations were found in multiple samples and because non-serine translations decreased considerably when analysing the data with the pseudo-haploid WO-1 annotation instead of the diploid SC5314 annotation. Third, we found similar levels of ambiguity for other rare leucine and serine codons which indicates that the observed mistranslation level corresponds to the general ribosomal mistranslation rate. This is further supported by observation of similar mistranslation rates in other yeasts. If the assumed 3-5% mistranslation of CUG codons by leucine were below the detection level of our approach, the background ambiguity we observed would be at an even higher level. In summary, our findings suggest that *C. albicans* does not decode CUG ambiguously. Phenotypic diversity is, therefore, not caused by CUG decoding unambiguity, a mechanism suggested to allow *C. albicans* to explore available ecological landscapes (Bezerra et al. 2013).

## Materials and Methods

### Growth and lysis of fungal cells

*Candida dubliniensis* CBS 7987 (CD36) and *Millerozyma acaciae* CBS 5656 (JCM 10732) were obtained from the CBS-KNAW culture collection of the Westerdijk Fungal Biodiversity Institute, Netherlands. The other strains are part of the yeast collection of the Institute of Microbiology of the University Medical Center Göttingen. *C. albicans* DSM70014, *C. dubliniensis* CBS 7987 and *Candida tropicalis* DSM 24507 were grown in YEPD medium (containing [% w/v]: bacto peptone 2.0; yeast extract 1.0; glucose 2.0) at 37°C. *M. acaciae* was grown in YEPD medium at 30°C. Cells were harvested and lysed exactly as described earlier (Mühlhausen et al. 2018). Proteins were resuspended in SDS sample buffer and resolved on 4-12 % gradient SDS-PAGE. MLST-typed (Odds et al. 2007) *C. albicans* isolates SC5314 (dST 52, clade 1), EU0006 (dST 1321, clade 2), EU0009 (dST299, clade 12) and EU0075 (dST124, clade 4) were subcultured three times on SAB agar after thawing. For production of yeast form cells, 50 ml Lee’s medium (Lee et al. 1975) with a pH of 4.5 were inoculated with a fresh colony and incubated at 30°C and 140 rpm in an orbital shaker overnight. This overnight culture was then diluted to an OD of 1.0 in 150 ml Lee’s pH 4.5 and grown at 30°C and 170 rpm in an orbital shaker. Hyphae form cells were produced starting from the above yeast overnight culture. Cells were treated identically as above except for using Lee’s medium with pH 6.5 and shaking at 37°C. Cells from 100 ml fungal culture were harvested during exponential phase (after ∼ 6 hours) by centrifugation at 4700x***g*** for 5 min. The pellets were washed in 10 ml fresh Lee’s medium (pH 4.5 or 6.5, respectively). Subsequently, 1 ml aliquots were taken from the cell suspension, centrifuged at 15,600x***g***, the supernatants were discarded and cells were shock frozen in liquid nitrogen and stored at −20°C. For SDS-PAGE, cell pellets were defrosted, 50 µl of loading buffer added, and samples vortexed for 10 sec. For mechanical cell lysis, glass beads were added and samples were passed twice for 20 sec in a FastPrep machine (speed setting 4.0). In between, samples were cooled on ice for 1 min. Proteins were resolved on 12% SDS-PAGE.

### Genome assemblies and annotation

Genome annotations for *C. albicans* SC5314 A22 (assembly 22), *C. albicans* WO-1, *C. dubliniensis* CD36, and *C. tropicalis* MYA-3404 were obtained on 18/11/2018 from the Candida Genome Database (Skrzypek et al. 2017). The *M. acaciae* JCM 10732 genome assembly was downloaded from GenBank with the accession BCKO01000000 (Shen et al. 2018). *M. acaciae* JCM 10732 genes were predicted with AUGUSTUS (Stanke and Waack 2003) using the parameter “genemodel=complete”, the gene feature set of *Candida albicans* and the standard codon translation table.

### Dilution of *M. acaciae* samples with *E. coli*

Protein concentration of *M. acaciae* and *E. coli* cell lysates was determined using the Biuret method. As starting sample, a mixture of 90% *M. acaciae* and 10% *E. coli* was produced, subsequently an aliquot of this sample mixed with an identical volume of the *M. acaciae* cell lysate, and the latter process of diluting the *E. coli* concentration by half repeated several times. This way, pipetting errors were reduced as much as possible.

### Mass spectrometric sequencing

SDS-PAGE-separated protein samples were processed as described by Shevchenko *et al.* (Shevchenko et al. 1996). The resuspended peptides in sample loading buffer (2% acetonitrile and 0.05% trifluoroacetic acid) were separated and analysed on an UltiMate 3000 RSLCnano HPLC system (Thermo Fisher Scientific) coupled online to either a Q Exactive HF or an Orbitrap Fusion mass spectrometer (Thermo Fisher Scientific). Firstly, the peptides were desalted on a reverse phase C18 pre-column (Dionex 5 mm long, 0.3 mm inner diameter) for 3 minutes. After 3 minutes the precolumn was switched online with the analytical column (30 cm long, 75 μm inner diameter) prepared in-house using ReproSil-Pur C18 AQ 1.9 μm reversed phase resin (Dr. Maisch GmbH). The peptides were separated with a linear gradient of 5–45% buffer (80% acetonitrile and 0.1% formic acid) at a flow rate of 300 nl/min (with back pressure 500 bars) over 58 min gradient time. The pre- and main column temperatures were maintained at 50°C. In the Q Exactive Plus the MS data were acquired by scanning the precursors in mass range from 350 to 1600 m/z at a resolution of 70,000 at m/z 200. Top 20 precursor ions were chosen for MS2 by using data-dependent acquisition (DDA) mode at a resolution of 17,500 at m/z 200 with maximum IT of 50 ms. In the Q Exactive HF the MS data were acquired by scanning the precursors in mass range from 350 to 1600 m/z at a resolution of 60,000 at m/z 200. Top 30 precursor ions were chosen for MS2 by DDA mode at a resolution of 15,000 at m/z 200 with maximum IT of 50 ms. Data were measured on Q Exactive HF instrument except for *M. acaciae* experiments which were measured on Orbitrap Fusion.

### Mass spectrometry data analysis

Data analysis and search were performed using MaxQuant v.1.6.0.1 (samples Calbicans_WO1_1, Calbicans_SC5314_1, *C. dubliniensis, C. glabrata* and *C. tropicalis*) and MaxQuant v.1.6.5.0 (all other samples) as search engine with a global and a peptide-level 1% FDR. To obtain peptide mappings free of codon-translation bias, 19 replicates for each genome annotation were generated with the codon translated as different amino acid in each replicate (translation as isoleucine being omitted as leucine and isoleucine are indistinguishable through MSMS). To reduce database size and redundancy, predicted proteins were split at lysine and arginine residues into peptides resembling trypsin proteolysis. Peptides containing the respective codons were fused together with the two subsequent peptides so that codon-containing fragments can be detected with up to two missed cleavages. The remaining peptides were fused back together as long as they formed consecutive blocks. Duplicate peptides (originating from peptide-blocks without the codon of interest) were removed. Search parameters for searching the precursor and fragment ion masses against the databases were as described in Mühlhausen *et al.* (Mühlhausen et al. 2018). To claim codon translations with high confidence, we determined whether the respective codon positions are supported by b- and y-type fragment ions at both sides allowing determination of the amino acids’ mass. Only those positions were regarded as fully supported by the data.

## Data access

The mass spectrometry data from this study have been submitted to the ProteomeXchange Consortium (http://proteomecentral.proteomexchange.org) via the PRIDE (Vizcaíno et al. 2016) partner repository with the data set identifier xxxxx.

## Acknowledgements

We thank Alexander Stein for providing the *E. coli* lysate.

## Author contributions

MK initiated the study. SM performed MS data analyses. HDS, PM, and PS prepared experimental samples. UP performed MS experiments. TP, MW and OB supervised sample preparation and were involved in data interpretation. HU was involved in MS data interpretation. MK assembled, aligned and analysed tRNA sequences. MK wrote the manuscript with support from SM and OB.

